# A neuron, microglia, and astrocyte triple coculture model to study Alzheimer disease

**DOI:** 10.1101/2021.12.13.472367

**Authors:** Celia Luchena, Jone Zuazo-Ibarra, Jorge Valero, Carlos Matute, Elena Alberdi, Estibaliz Capetillo-Zarate

**Affiliations:** Achucarro Basque Center for Neuroscience. Leioa, Spain; Facultad de Medicina y Enfermería, Universidad del País Vasco. Leioa, Spain; CIBERNED, Centro de Investigación Biomédica en Red Enfermedades Neurodegenerativas. Spain; IKERBASQUE, Basque Foundation for Science. Bilbao, Spain; Institute of Neurosciences of Castilla y León, University of Salamanca, Salamanca, Spain, and Institute for Biomedical Research of Salamanca, Salamanca, Spain

**Keywords:** neuron, astrocyte, microglia, in vitro, co-culture, Alzheimer, synapse

## Abstract

Glial cells are essential to understand Alzheimer’s disease (AD) progression, given their role in neuroinflammation and neurodegeneration. There is a need for reliable and easy to manipulate models that allow studying the mechanisms behind neuron and glia communication. Currently available models such as cocultures require complex methodologies and/or might not be affordable for all laboratories. With this in mind, we aimed to establish a straightforward *in vitro* setting with neurons and glial cells to study AD.

We generated a triple co-culture with neurons, microglia and astrocytes. Immunofluorescence, western blot and ELISA techniques were used to characterize the effects of oligomeric Aβ (oAβ) in this model.

We found that, in the triple co-culture, microglia increased the expression of anti-inflammatory markers Arginase I and TGF-β1, and reduced pro-inflammatory iNOS and IL-1β, compared with microglia alone. Astrocytes reduced expression of pro-inflammatory A1 markers AMIGO2 and C3, and displayed a ramified morphology resembling physiological conditions. Lastly, neurons increased post-synaptic markers, and developed more and longer branches than in individual primary cultures. Addition of oAβ in the triple coculture reduced synaptic markers and increased microglial activation, which are hallmarks of AD.

Consequently, we developed a reliable model, where cells better resemble physiological conditions: microglia are less inflammatory, astrocytes are less reactive and neurons display a more mature morphology than in individual primary cultures. Moreover, we were able to recapitulate Aβ-induced synaptic loss and inflammation. This model emerges as a powerful tool to study neurodegeneration and inflammation in the context of AD and other neurodegenerative diseases.

**Table of content image:** 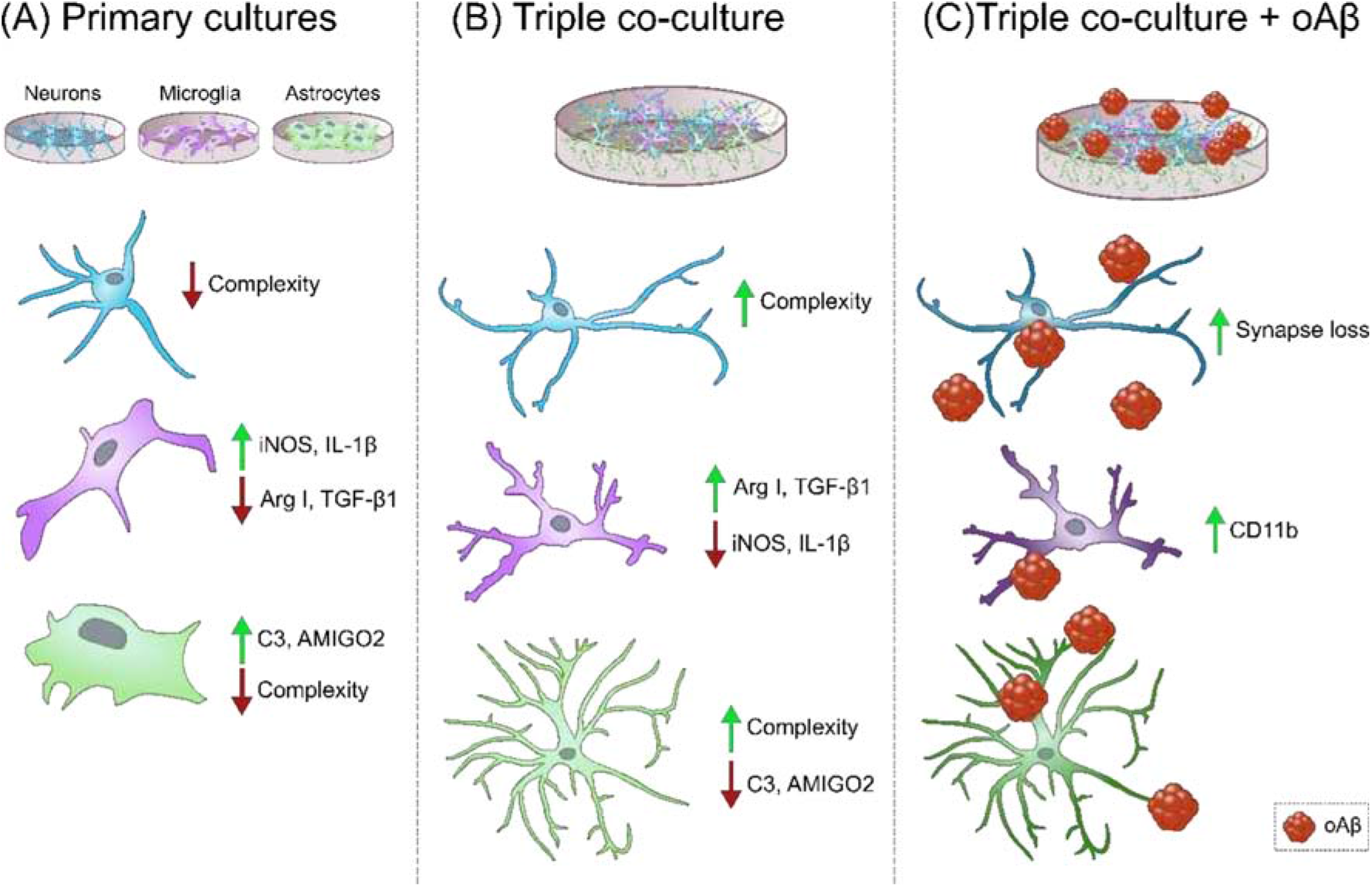

**Main points:**

- In our model, microglia and astrocytes are less reactive, and neurons have a more mature morphology than in primary cultures.
- oAβ reduced synaptic markers and increased microglial activation.
- This triple co-culture is a reliable tool to study neurodegeneration and gliosis *in vitro*.

## 1. Introduction

Alzheimer’s disease (AD) is a neurodegenerative disease characterized, among other hallmarks, by amyloid-β (Aβ) peptide aggregation and accumulation, synapse loss and inflammation (Castellani *et al*., 2010). AD is a multifactorial disease, and thus, understanding the relationship between glial cells and neurons is essential to comprehend the highly complex mechanisms triggered by the accumulation of Aβ.

The use of animal models of AD has demonstrated that targeting glial cells can ameliorate the disease progression (Furman *et al*., 2012; Czirr *et al*., 2017). While AD animal models allow researchers to study the complexity of neuropathology and inflammation, mechanistic studies together with cell-to-cell communication analyses are limited due to the inherent large amount of variables present in animal models. *In vivo* models might also require long time courses and high financial efforts. Moreover, high number of animals might be required. On the other hand, widely used *in vitro* models such as individual primary cultures, immortalized cell lines and, more recently, human-derived cells are more susceptible to manipulation, time scales are shorter, and the amount of animals required is smaller. However, they lack complexity. Different co-culture models have been developed in recent years to study the interaction between glial cells and neurons. To study cell communication via secreted components, conditioned media have been extensively used. For example, Aβ-stimulated conditioned media from microglia was discovered to stimulate neurotoxicity via secreted TNFα and NMDA receptor stimulation in mouse primary neurons (Floden *et al*., 2005). Secreted factors from astrocytes can also affect neuronal function. Christopherson *et al*. found that conditioned media from immature astrocytes contains thrombospondins-1 and −2, which promote synaptogenesis in retinal ganglion cells (Christopherson *et al*., 2005). In another study, conditioned media from LPS-activated microglia containing IL-1α, TNF and C1q, induced A1 phenotype in astrocytes, which reduced their capacity to promote neuronal survival and synaptogenesis (Liddelow *et al*., 2017). Other researchers have developed co-culture models where the two cell populations share the medium but are not in contact with each other. For instance, Du *et al*. established an iPSC-derived neuron-astrocyte co-culture using transwell inserts, and found that astrocytes rescued mitochondrial dysfunction in dopaminergic neurons treated with mitochondrial toxins (Du *et al*., 2018). Another advantage of co-cultures is that they can mitigate the drastic transcriptomic modifications that cells, especially microglia, undergo due to manipulation and the modification of their environment. Transcriptomic profiling of human and mouse microglia has shown significant downregulation of genes related to immune signaling and brain development, and upregulation of genes related to inflammation and stress responses in *in vitro* conditions, compared to *ex vivo* (Gosselin *et al*., 2017). Microglia *in vitro* and in the brain also have different responses to Aβ. Murine primary microglia display rapid and substantial transcriptional changes in response to Aβ, while *in vivo* microglia do not display the same gene expression changes, suggesting that primary microglia poorly recapitulate *in vivo* conditions (McFarland *et al*., 2021). Nevertheless, studies have shown that culturing microglia together with neurons can mitigate this problem. iPSC-derived microglia co-cultured with neurons express key microglia-specific markers, display dynamic ramifications, and have phagocytic capacity. They also upregulate homeostatic genes and promote a more anti-inflammatory cytokine profile than monocultures (Haenseler *et al*., 2017). In recent years, more complex co-cultures have been developed using three cell populations to study intercellular communication. Park *et al*. developed a human triculture model using human neurons, astrocytes, and microglia in a 3D microfluidic platform. With this model, they were able to recapitulate AD features such as Aβ aggregation, tau phosphorylation, and microglial recruitment. They were also able to demonstrate neurotoxic effects derived from neuron-glia interactions (Park *et al*., 2018). In a different study, a tri-culture system containing hPSC-derived neurons, microglia and astrocytes was used to study complement component C3, and found a reciprocal signaling between microglia and astrocytes that resulted in increased C3 in AD conditions (Guttikonda *et al*., 2021). However, these models might involve complex methodology and/or might not be so affordable. In this study, we developed and characterized a murine triple co-culture model with neurons, astrocytes and microglia that can be used to study cellular communication in the context of Aβ pathology and neuroinflammation. It is an inexpensive and straightforward model that better resembles physiological conditions compared to individual primary cultures, and it represents a more reliable *in vitro* model for mechanistic studies of neurodegenerative diseases.

## 2. Materials and methods

### 2.1. Animals

Sprague-Dawley wild-type rats of both genders were used to obtain cell cultures and organotypic slices. All animal experiments were compliant with the requirements of the Animal Ethics and Welfare Committee of the University of the Basque Country (UPV/EHU), following the Spanish Real Order 53/2013 and the European Communities Council Directive of September 22^nd^ 2010 (2010/63/EU).

### 2.2. Cortical neuron culture procedures

Neurons were obtained from the cortical lobes of E18/E19 Sprague-Dawley rat embryos as previously described (Larm *et al*., 1996; Ibarretxe *et al*., 2006), with modifications. Cortical lobes were digested in 0.2% trypsin and 0.02% deoxyribonuclease I in Hanks’ Balanced Salts, no calcium, no magnesium (HBSS) (Sigma-Aldrich, MO, USA) for 5 min at 37°C. The enzymatic digestion was stopped by adding Neurobasal medium containing 2% B27 (Gibco, MA, USA) (Brewer *et al*., 1993), 1% Penicillin-Streptomycin (Lonza, Switzerland) and 0.5 mM L-glutamine, plus 10% fetal bovine serum (FBS) (Gibco, MA, USA). After incubating 10 min at 37°C, cells were centrifuged for 5 min at 200 g, and the resulting pellet was resuspended in B27 Neurobasal medium plus 10% FBS. Cells were mechanically dissociated by trituration (20 strokes with 21, 23 and 25-gauge needles), filtered through a 41 μm nylon mesh and seeded onto poly-D-lysine-coated (PDL) (Sigma-Aldrich, MO, USA) plates as needed. The medium was replaced by serum-free B27 Neurobasal 24 hours later. Cultures were maintained at 37°C and 5% CO_2_ for eight to nine days *in vitro* (DIV) before use.

### 2.3. Cortical glial culture procedures

Primary cultures of mixed glial cells (source cultures) were prepared from the cortical lobes of P0/P1 Sprague-Dawley rat pups as described elsewhere (McCarthy and De Vellis, 1978). Cortical lobes were incubated with 0.2% trypsin and 0.01% deoxyribonuclease I in HBSS for 15 min at 37°C. The digestion was stopped adding Iscove’s Modified Dubecco’s Medium (IMDM) (Gibco, MA, USA) with 1% Antibiotic-Antimycotic (Gibco, MA, USA), plus 10% HyClone™ FBS (Cytiva, MA, USA). Cells were centrifuged for 6 min at 1200 rpm (equivalent to approx. 250 g), and the pellet was resuspended in IMDM plus 10% HyClone™ FBS. This pellet was mechanically dissociated by trituration (20 strokes with 21 and 23-gauge needles) and recentrifuged. Cells were resuspended in IMDM plus 10% HyClone™ FBS and plated onto PDL-coated 75 cm^2^ flasks (about four cortices per flask). One day later, the medium was replaced by fresh IMDM plus 10% HyClone™ FBS for primary astrocytes, or by Dulbecco’s Modified Eagle Medium high glucose, pyruvate (DMEM) (Gibco, MA, USA) with 1% Penicillin-Streptomycin (Lonza, Switzerland) plus 10% FBS for primary microglia. Cultures were maintained at 37°C and 5% CO_2_ for one week to collect astrocytes, or for two weeks to collect microglia.

Primary astrocytes were obtained from source mixed glial cultures as described previously (McCarthy and De Vellis, 1980; Alberdi *et al*., 2013). After one week of incubation, medium was removed, and flasks were washed twice with HBSS. Cells were trypsinized by incubating with Trypsin-EDTA 0.05% (Gibco, MA, USA) for 5 min at 37°C. IMDM plus 10% HyClone™ FBS was added to stop the enzymatic reaction and cells were centrifuged for 5 min at 1500 rpm (equivalent to approx. 300 g). The cell pellet was resuspended in IMDM plus 10% HyClone™ FBS and astrocytes were plated onto PDL-coated plates as needed. The medium was replaced by serum-free B27 Neurobasal 24 hours after plating. Cultures were maintained at 37°C and 5% CO_2_.

Primary microglia were also obtained from source mixed glial cultures as previously described (Paresce *et al*., 1997; Majumdar *et al*., 2011). After growing for two weeks, microglia were harvested by orbital shaking for 1 h at 400 rpm in DMEM plus 10% FBS. Cells were centrifuged for 6 min at 1200 rpm (equivalent to approx. 250 g) and plated onto PDL-coated plates as needed. The medium was replaced by serum-free B27 Neurobasal 24 hours after plating. Cultures were maintained at 37°C and 5% CO_2_.

### 2.4. Triple co-culture procedures

To obtain all possible combinations of neuron-glia co-cultures (neuron-astrocyte, neuron-microglia and neuron-astrocyte-microglia), primary cortical cell cultures were prepared as stated above. Cells were plated at different ratios as it has been described (Hernangómez *et al*., 2012; Correa *et al*., 2013) to find the adequate conditions. The optimal conditions are described in Figure 1A. The co-cultures were started by plating astrocytes onto PDL-coated plates. When a monolayer of astrocytes was obtained three days later, neurons were plated in a proportion of five neurons to two astrocytes. After 24 h, the medium was replaced by serum-free B27 Neurobasal, and antimitotic agents 5-Fluoro-2′-deoxyuridine and Uridine (FdU/U) (Sigma-Aldrich, MO, USA) were added at a concentration of 10μM to control glial proliferation. Cells were incubated at 37°C and 5% CO_2_, and more FdU/U could be added every two days for further control of proliferation. After six days, microglia was added to the cultures in a proportion of one microglia to five neurons. Cultures were ready to process for protein analysis two days later. For experiments with oligomeric Aβ (oAβ), a treatment of 3 μM oAβ for 24 h was used. The co-cultures were kept for a total of 13 DIV at 37°C and 5% CO_2_ in serum-free B27 Neurobasal.

**Figure 1.**
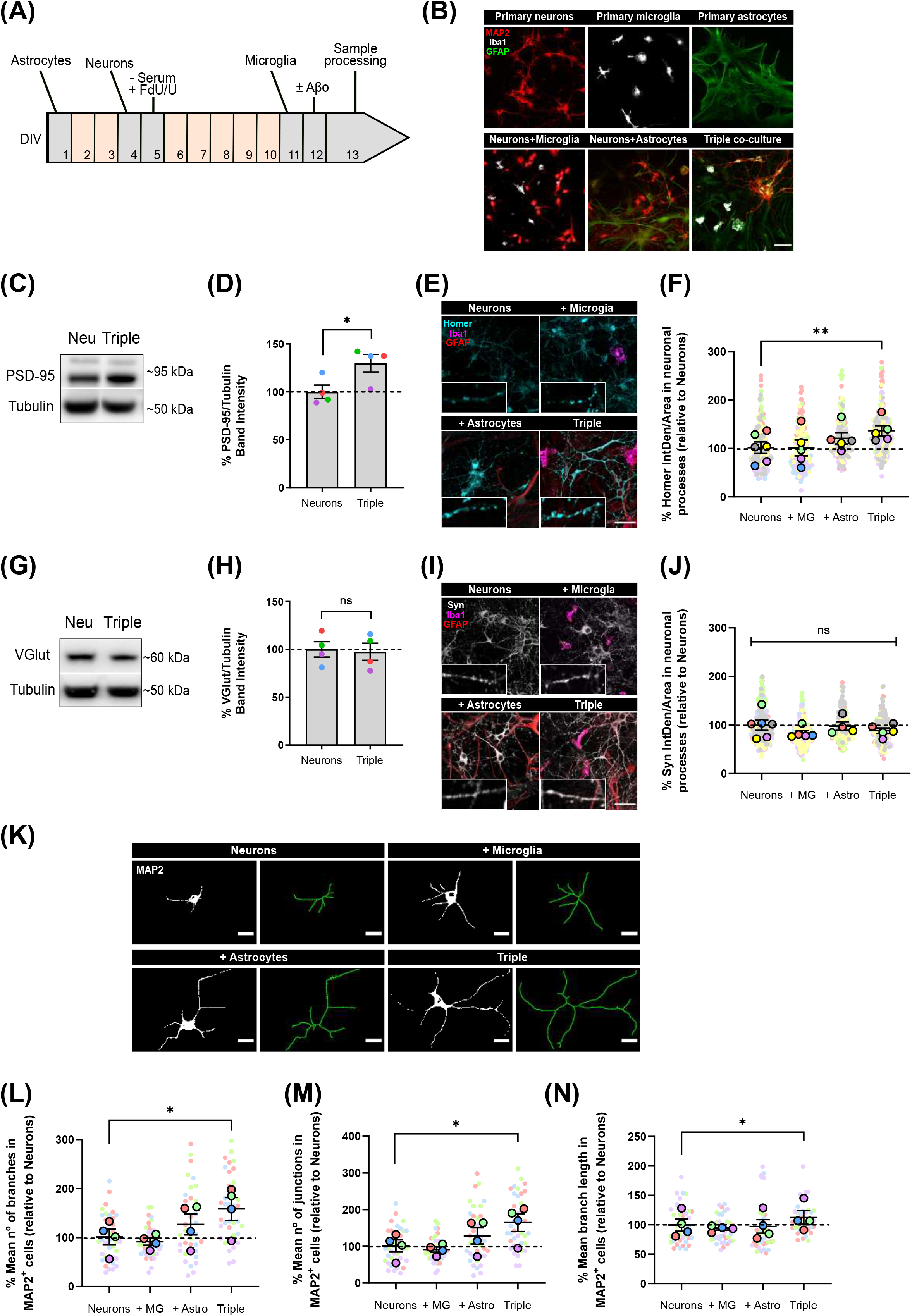
Neurons increased the expression of post-synaptic markers PSD-95 and Homer1, and developed more and longer branches when cocultured with microglia and astrocytes. (A) Protocol diagram for the *in vitro* triple co-culture model and (B) representative images. Cultures started with a monolayer of astrocytes. Neurons were plated three days later (ratio: five neurons to two astrocytes), and microglia were added seven days after that (ratio: one microglia to five neurons). Cultures were maintained for 13 DIV in total. Scale bar = 50 μm. (C) Representative western blot image of the quantification of PSD-95 in total lysates. (D) Post-synaptic PSD-95 significantly increased (30.0 ± 9.1%; p = 0.0408; n = 4) in the triple co-culture compared with neurons alone by western blot. (E) Representative images of Homer1 immunofluorescence in neuronal processes. Scale bar = 40 μm. (F) Post-synaptic Homer1 increased (34.5 ± 10.5%; p = 0.0093; n = 6) in the triple coculture, compared with neurons alone. (G) Representative western blot image of the quantification of VGlut1 in total lysates. (H) VGlu1 did not change in the triple co-culture compared with neurons alone by western blot. (I) Representative images of Synaptophysin immunofluorescence in neuronal processes. Scale bar = 40 μm. (J) Pre-synaptic Synaptophysin did not change in any condition. (K) Representative images of the quantification of cellular morphology using the neuronal marker MAP2 and the function Skeletonize in FIJI software. Scale bar = 20 μm. (L) Neurons displayed significantly higher number of branches (56.7 ± 23.3%; p = 0.0290; n = 4), (M) junctions (62.6 ± 24.2%; p = 0.0190; n = 4) and (N) branch length (12.8 ± 11.5%; p = 0.0431; n = 4) in the triple co-culture. Each bar represents the mean ± SEM. * p<0.05; ** p<0.01; ^ns^ not significant. DIV= days *in vitro*; Neu = neurons; MG = microglia; Astro = astrocytes; Syn = synaptophysin.

### 2.5. Conditioned media experiments

To obtain conditioned media, primary neurons, astrocytes and microglia were seeded separately at a density of 1×10^6^ cells in Petri dishes (60 mm in diameter). One day later, medium was replaced by serum-free B27 Neurobasal, and cells were incubated for three days at 37°C and 5% CO_2_. Supernatants were collected, stored at −80°C and filtered with a 22-μm nylon mesh before use.

### 2.6. Preparation of soluble oligomeric Aβ_1-42_

Synthetic Aβ oligomers were prepared as previously reported (Dahlgren *et al*.,2002; Alberdi *et al*., 2010). Briefly, Aβ_1-42_ (Bachem, Switzerland) was dissolved in hexafluoroisopropanol (HFIP) (Sigma-Aldrich, MO, USA) to an initial concentration of 1 mM. It was distributed into aliquots, and HFIP was removed under vacuum in a speed vac system before the peptide film was stored dried at −80°C. For the aggregation protocol, the peptide was resuspended in anhydrous dimethylsulfoxide (DMSO) (Invitrogen, MA, USA) to a concentration of 5 mM. Hams F-12 pH 7.4 (Biowest, France) was added to reach the final concentration of 100 μM. Oligomers were obtained after incubating at 4°C for 24 h.

### 2.7. Organotypic Hippocampal Slice Culture

Hippocampal slice cultures were obtained from P5/P6 Sprague-Dawley rat pups and prepared as previously described (Stoppini *et al*., 1991; Ortiz-Sanz *et al*., 2020) with modifications. Brains were extracted in ice cold HBSS, and chopped into 400 μm thick slices using a vibratome (VT 1200S, Leica, Germany). Hippocampi were isolated from intact slices under a dissection microscope, and transferred onto Millicell culture inserts with 30 mm in diameter (EMD Millipore, MA, USA) with fresh Neurobasal medium containing 25% horse serum (HS) (Gibco, MA, USA), 22% HBSS, 1% D-glucose 550 mM, 1% L-glutamine 200 mM and 1% Antibiotic-Antimycotic below the membranes. Tissue slices were maintained at 37°C and 5% CO_2_ and the medium was renewed every two days. Experiments were performed at 14-15 DIV. One day before treatment, the medium was replaced by Neurobasal medium supplemented with 1% HS, 25% HBSS, 1.8% D-glucose 550 mM, 1% L-glutamine 200 mM and 1% Antibiotic-Antimycotic. For experiments with Aβ, slices were treated with 3 μM oAβ for 24 h.

### 2.8. Western blotting

Cell lysates were prepared from cultures using a cell scraper (Corning, NY, USA) and Pierce™ RIPA buffer, with Halt™Protease & Phosphatase Inhibitor Cocktail (Thermo Fisher Scientific, MA, USA) and 0.5 M ethylenediamine tetraacetic acid solution (EDTA). Protein concentrations were determined by DC Protein Assay. Samples were diluted in 2X sodium dodecyl sulfate (SDS) sample buffer and boiled at 95°C for 5 min before use. Equal amounts of protein lysates were loaded onto Bolt™ 4 - 12% Bis-Tris mini protein gels, using Bolt™ MES SDS Running buffer (Invitrogen, MA, USA) for electrophoresis, and transferred to iBlot™ 2 PVDF membranes (Invitrogen, MA, USA), followed by immunoblotting. Membranes were blocked in 1X tris-buffered saline (TBS) (20 mM Tris, 137 mM NaCl in double distilled water (ddH_2_O), pH 7.4), with 0.05% Tween-20 (TBS-T) (Acros organics, Belgium) and 5% fat-free milk for 1 h at room temperature (RT). Proteins were detected by incubation overnight with specific primary antibodies against PSD-95 (1:1000, ab18258, abcam, UK), VGlut1 (1:750, 135511, Synaptic Systems, Germany), Alpha tubulin (1:2000, ab7291, abcam, UK), MRC1 (1:1000, ab64693, abcam, UK), Arginase I (1:200, sc-166920, Santa Cruz Biotechnology, CA, USA), Iba1 (1:500, 016-20001, Fujifilm Wako, Japan), CD11b (1:500, ab128797, abcam, UK) and GAPDH (1:2000, MAB374, Sigma-Aldrich, MO, USA). Membranes were incubated with HRP-linked secondary antibodies anti-mouse IgG (1:5000, A6782, Sigma-Aldrich, MO, USA) and anti-rabbit IgG (1:5000, 7074S, Cell Signaling Technology, MA, USA). Bands were detected with a ChemiDoc™ MP Imaging System (Bio-Rad, CA, USA), and the band intensities were quantified using Bio-Rad Image Lab^®^ software. All protein intensities were divided by the corresponding tubulin or GAPDH measurement for normalization, unless otherwise stated.

### 2.9. ELISA Assays

Pro-inflammatory cytokine Interleukin-1β (IL-1β) present in supernatants was measured using an appropriate ELISA kit (ab100767, abcam, UK), following manufacturer’s instructions. Supernatant samples from cell cultures (density of 1×10^5^ - 1×10^6^ cells) were used undiluted. In brief, 100 μl of standards and samples were added into a 96-well plate pre-coated for rat IL-1β and incubated for 2.5 h at RT with gentle shaking. The plate was washed with washing solution, and 100 μl of Biotinylated IL-1β detection antibody were added to each well. After incubating for 1 h at RT with gentle shaking, the plate was washed again, and 100 μl of HRP-Streptavidin solution were added to each well. The plate was incubated for 45 min at RT with gentle shaking and washed before adding 100 μl of TMB One-Step substrate reagent to each well. After a 30 min incubation at RT with gentle shaking in darkness, 50 μl of Stop solution were added and the plate was read immediately in a fluorimeter at 450 nm. The standard curve was obtained plotting the standard concentration (pg/ml) on the x-axis and absorbance on the y-axis. IL-1β concentrations in each sample were calculated using the equation of the best-fit straight line through the standard points.

Levels of anti-inflammatory cytokine Transforming growth factor-β1 (TGF-β1) in supernatants were measured by an ELISA kit (ab119558, abcam, UK), following manufacturer’s instructions. Amicon®Ultracel®3K centrifuge filters (EMD Millipore, MA, USA) were used to concentrate the amount of proteins present in the cell culture supernatants (density of 1×10^5^ - 1×10^6^ cells), due to low levels of TGF-β1. For the ELISA assay, 20 μl of all samples were prediluted with 180 μl of Assay buffer plus 20 μl of 1N HCl for 1h, and then neutralized with 20 μl 1N NaOH. 100 μl of standards and samples were added into a 96-well plate precoated for rat TGF-β1 and incubated for 2 h at RT with gentle shaking. The plate was washed with washing solution, and 100 μl of Biotin-Conjugated Antibody were added to each well. After incubating for 1 h at RT with gentle shaking, the plate was washed again, and 100 μl of Streptavidin-HRP solution were added to each well. The plate was incubated for 30 min at RT with gentle shaking and washed before adding 100 μl of TMB Substrate Solution to each well. After a 30 min incubation at RT with gentle shaking in darkness, 100 μl of Stop solution were added and the plate was read immediately in a fluorimeter at 450 nm. The standard curve was obtained plotting the standard concentration (pg/ml) on the x-axis and absorbance on the y-axis. TGF-β1 concentration in each sample was calculated using the equation of the best-fit straight line through the standard points.

### 2.10. Inmunofluorescence

Cell culture coverslips were fixed in 4% PFA for 10 min, and washed twice with 1X TBS. Then, they were permeabilized and blocked in 1X TBS with 1% BSA, 0.5% normal goat serum (NGS) (Palex Medical, Spain) and 0.1% saponin for 1 h. Coverslips were incubated overnight at 4°C with primary antibodies against MAP2 (1:1000, #M9942, Sigma-Aldrich, MO, USA), Iba1 (1:500, #234004, Sinaptic Systems, Germany), GFAP (1:1000, #MAB3402, Sigma-Aldrich, MO, USA), Synaptophysin (1:1000, #837101, BioLegend, CA, USA), Homer1 (1:500, #160003, Synaptic Systems, Germany), iNOS (1:200, #610329, BD Biosciences, NJ, USA), CD11b (1:500, #ab128797, abcam, UK), C3 (1:100, #ab11887, abcam, UK), PSD95 (1:500, #ab18258, abcam, UK), AMIGO2 (1:250, #sc-373699, Santa Cruz Biotechnology, CA, USA) and GFAP (1:4000, #Z0334, Dako, CA, USA). Samples were washed with TBS-T and incubated with Alexa fluorophore-conjugated secondary antibodies (1:500, Invitrogen, MA, USA) for 1h at RT. Cell nuclei were stained by incubating with 5 μg/ml DAPI (Sigma-Aldrich, MO, USA) for 15 min at RT, and samples were mounted using Fluoromont-G (SouthernBiotech, AL, USA).

Organotypic hippocampal slices were fixed in 4% PFA for 40 min, washed twice with 1X phosphate-buffered saline (PBS) (125 mM NaCl, 19 mM Na_2_HPO_4_, 8 mM KH_2_PO_4_ in ddH_2_O, pH 7.4), and the membrane was trimmed around each slice with a scalpel for better handling. Samples were permeabilized and blocked for 3 h at RT in 1X PBS with 5% NGS and 0.5% Triton-X. Then, slices were incubated overnight at 4°C in 1X PBS with 1% NGS and 0.1% Triton-X with primary antibodies against Synaptophysin (1:1000, #837101, BioLegend, CA, USA), PSD95 (1:500, #ab18258, abcam, UK) and CD11b (1:500, #MCA275R, Bio-Rad, CA, USA). Samples were washed with 1X PBS and incubated with Alexa fluorophore-conjugated secondary antibodies (1:500, Invitrogen, MA, USA) for 2h at RT. Slices were mounted using a bridge mounting technique (Yoon *et al*., 2010). Two 22 mm x 22 mm coverslips were glued to each end of a glass slide. Organotypic slices were mounted using Fluoromont-G, and a 40 mm x 22 mm coverslip was placed to bridge the gap without smashing the samples.

### 2.11. Confocal microscopy and image processing

Images were acquired using Leica TCS STED CW SP8X confocal microscope (Leica, Germany). In experiments with multiple fluorophores, channels were scanned sequentially to avoid crosstalk. The same settings were applied to all images within the same experiment. All analyses were carried out using open source ImageJ/Fiji software.

Synaptic markers were quantified in neuritic segments of primary neurons. In each condition, images (size 184×184 μm, resolution 5.55 pixels/micron) were acquired on random fields, and neuritic segments (20 μm in length) were selected from areas where a single process could be outlined. After the background was subtracted and a threshold was applied, integrated density of each neuritic segment was quantified. For each condition, individual segments were grouped and averaged per field, and all the per field averages were used to calculate the group mean and standard deviation for each condition and experiment. A macro was developed in ImageJ/Fiji software to automate the analysis.

For neuronal morphology analyses, MAP2^+^ positive cells were used. Single neurons were selected in areas where a single cell could be identified. Images were converted to binary and then skeletonized. The function Summarize skeleton in ImageJ/Fiji software was used to obtain the average branch length, the number of branches and the number of junctions in each neuron. The values of each individual cell were used to obtain the group mean and standard deviation for each condition and experiment.

To quantify the expression levels of proteins CD11b, iNOS and C3 in the cell cultures, the specific markers MAP2, GFAP and Iba1 were used to identify and define the area occupied by neurons, astrocytes and microglia, respectively. Then, the integrated density of each target protein was measured inside that delimited area. The integrated density of individual cells was averaged and divided by the total number of cells to obtain the integrated density per cell group mean and standard deviation for each condition and experiment.

To study astrocytic morphology in vitro, GFAP was used to stain astrocytes. They were selected in areas where a single astrocyte could be identified. After images of individual cells were transformed into binary, the tool Outline was used to automatically draw the cell shape. Morphology parameters like density (number of pixels of foreground color divided by the total number of pixels in the convex hull), span ratio (a measure of shape, as ratio of major and minor axes for the convex hull) and circularity were quantified using the Fractal Analysis FracLac plugin available in ImageJ/Fiji software as previously described (Young and Morrison, 2018). The values from individual cells were averaged to obtain the group mean and standard deviation for each condition and experiment.

For synaptic puncta colocalization studies, random fields in hippocampal organotypic slices were imaged with the 63X objective and 4X zoom in Leica TCS STED CW SP8X confocal microscope. Each image (size 34×34 μm, resolution 14.22 pixels/micron) was deconvoluted to better distinguish synaptic puncta using Huygens Professional software (Scientific Volume Imaging, The Netherlands). The number of pre-synaptic, post-synaptic and colocalized puncta (spots where the two synaptic channels overlapped) was determined for each image. The values from each image were averaged to obtain the group mean and standard deviation for each condition and experiment. A macro was developed in ImageJ/Fiji software to automate the analysis.

### 2.12. Statistical analysis

GraphPad Prism 8 (GraphPad Software, CA, USA) was used to perform all statistical analyses. Data were presented as mean ± SEM (standard error of the mean) and scaled so that the average value for the corresponding control was 100%. Each independent experiment was coded with a different color. All data sets were tested for normality and homoscedasticity. Two-tailed paired Student’s t test was used to compare two experimental groups. Comparisons between more than two groups were performed by paired one-way ANOVA, followed by *post hoc* Tukey test. If one or more data points were missing, mixed-effect model was used to perform the analysis. Statistical significance was represented as p<0.05 (*), p<0.01 (**), and p<0.001 (***).

## 3. Results

### 3.1. Neurons increase post-synaptic markers and develop a more complex morphology when co-cultured with glial cells

In order to establish an *in vitro* protocol where we could study neuron-glia communication and recapitulate neurodegeneration features such as synaptic loss, we optimized an easy and straightforward murine triple co-culture model (Figure 1A, B) with neurons, astrocytes and microglia in all possible combinations. We characterized each cell type in the co-culture and compared it with primary cultures.

Synapse formation and stabilization is promoted *in vitro* by astrocytes through secretion factors (Christopherson *et al*., 2005). Thus, we analyzed whether neurons would develop more synapses in the triple co-culture environment. To do so, we quantified pre- and post-synaptic markers in our co-cultures using western blotting (WB) and immunofluorescence (IF). We found that post-synaptic markers PSD-95 (30.0 ± 9.1%; p = 0.0408; n = 4) and Homer1 (34.5 ± 10.5%; p = 0.0093; n = 6) were increased in the triple co-culture compared with primary neurons alone (Figure 1C-F). Both microglia and astrocytes had to be present to observe this increase, because Homer1 was not altered in neuron-microglia or neuron-astrocyte co-cultures. Meanwhile, pre-synaptic markers VGlut1 (p = 0.8717; n = 4) and Synaptophysin (Syn) (p = 0.4053; n = 6) were not different in the triple co-culture compared to primary neurons (Figure 1G-J).

Studies carried out using dual co-cultures have suggested a change in neuronal morphology in the presence of microglia (Zhang and Fedoroff, 1996). Accordingly, we analyzed cellular morphology using the neuronal marker MAP2. In the triple co-culture, neurons displayed a higher number of branches (56.7 ± 23.3%; p = 0.0290; n = 4), as well as a higher number of junctions (62.6 ± 24.2%; p = 0.0190; n = 4). These branches were significantly longer (12.8 ± 11.5%; p = 0.0431; n = 4) compared with primary neurons alone (Figure 1K-N). It is worth noting that astrocytes alone showed a trend to increase the number of branches and junctions in neurons, although it was not statistically significant. In general, the conditions generated in the triple co-culture increased the number of post-synapses in neurons, as well as their cellular complexity.

### 3.2. Microglia display a less inflammatory phenotype when co-cultured with neurons and astrocytes

After observing that neurons exhibit different characteristics when glial cells were present, and given that microglial transcriptomic profiles are very dependent on the cellular environment (Schmid *et al*., 2009), we analyzed the possible phenotypic changes that microglia could undergo in our triple coculture model.

We used WB, IF and ELISA assays to quantify the expression of commonly used markers for pro- and anti-inflammatory microglial polarization. We found that the typically anti-inflammatory marker Arginase I (ArgI) displayed a ~ 8-fold increase in the triple co-culture (798.2 ± 142.7%; p = 0.0072; n = 4), as well as in the microglia-neuron co-culture (746.2 ± 147.3%; p = 0.0157; n = 4) compared with primary microglia alone (Figure 2A, B). No changes were found in the case of anti-inflammatory marker Mannose receptor C-type 1 (MRC1) (p = 0.0899; n = 4) (Figure 2A, C). We also found a ~ 2-fold increase of the antiinflammatory marker TGF-β1 in triple co-culture supernatants (132.6 ± 35.5%; p = 0.0067; n = 5) compared with primary microglia supernatants (Figure 2D). On the other hand, microglia decreased the secretion of pro-inflammatory cytokine IL-1β (34.2 ± 21.4%; p = 0.0059; n = 5) in the triple co-culture (Figure 2E). At the same time, there was a decrease of the pro-inflammatory marker iNOS in the triple co-culture (39.8 ± 7.6%; p = 0.0085; n = 4) and in the astrocyte-microglia co-culture (29.5 ± 11.3%; p = 0.0378; n = 4) compared with primary microglia alone (Figure 2F, G). No changes were observed in microglial activation marker CD11b (p = 0.3087; n = 4) (Figures 2F, H). In conclusion, microglia had a less inflammatory phenotype in the triple co-culture model compared with primary culture conditions, revealed by an increase of antiinflammatory markers ArgI and TGF-β1, and a decrease of pro-inflammatory markers IL-1β and iNOS.

**Figure 2.**
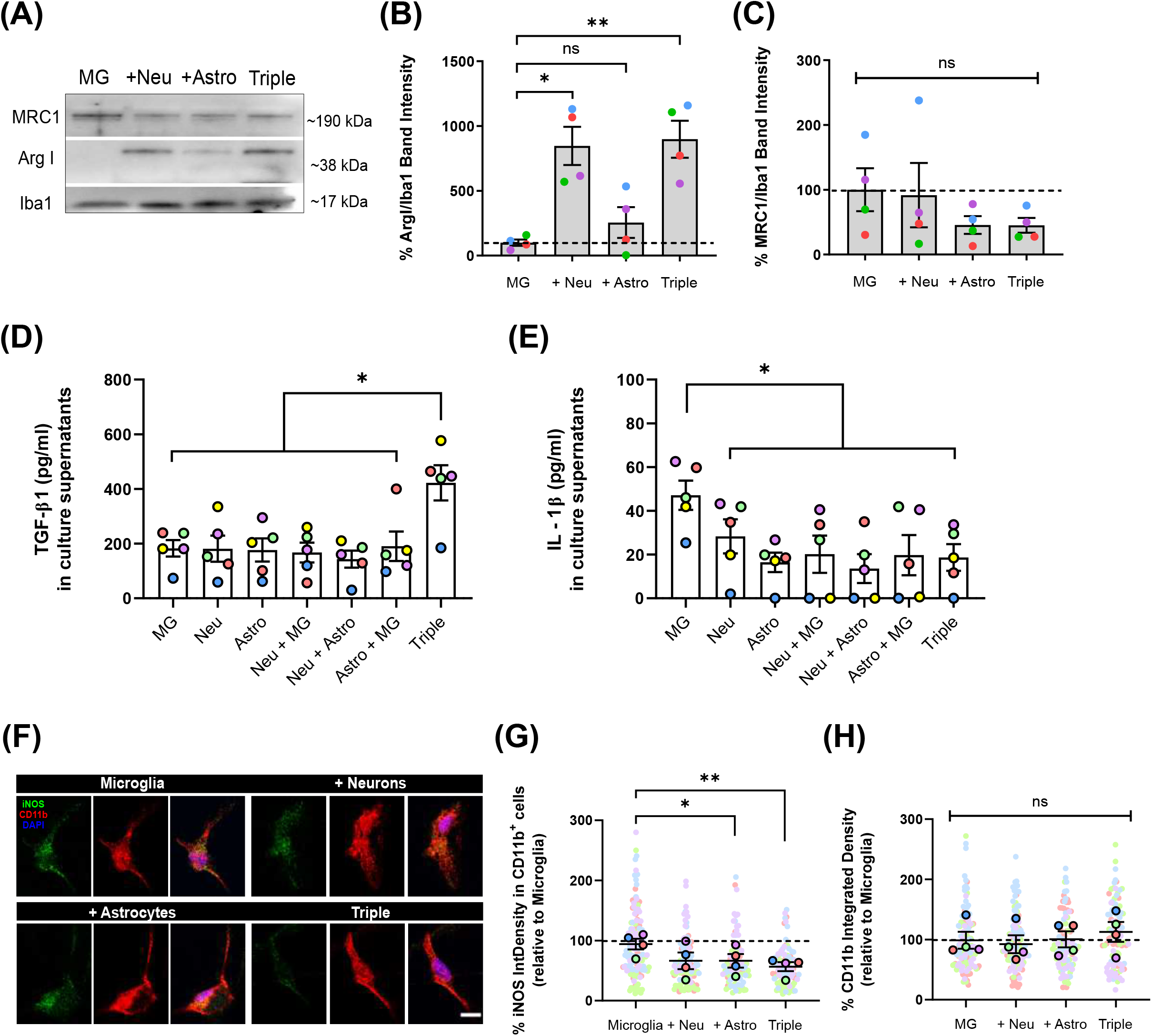
Microglia increased the expression of anti-inflammatory markers ArgI and TGF-β1, and decreased pro-inflammatory markers IL-1β and iNOS in the triple co-culture model. (A) Representative western blot image of microglial markers in total cell lysates. (B) Anti-inflammatory marker ArgI increased ~ 8-fold in the triple co-culture (798.2 ± 142.7%; p = 0.0072; n = 4) as well as in the microglia-neuron co-culture (746.2 ± 147.3%; p = 0.0157; n = 4) compared to primary microglia alone. Data was normalized using the marker Iba1 in order to selectively control for microglia. (C) Anti-inflammatory marker MRC1 did not change in any condition. (D) Anti-inflammatory cytokine TGF-β1 increased ~ 2-fold in the triple co-culture supernatants (132.6 ± 35.5%; p = 0.0067; n = 5) compared with primary microglia supernatants. (E) In culture supernatants, pro-inflammatory cytokine IL-1β decreased in the triple co-culture (34.2 ± 21.4%; p = 0.0059; n = 5) compared to primary microglia. (F) Quantification of microglial CD11b and pro-inflammatory marker iNOS. Scale bar = 10 μm. (G) The intensity of iNOS inside CD11b^+^ cells decreased in the triple co-culture (39.8 ± 7.6%; p = 0.0085; n = 4) and in the astrocyte-microglia co-culture (29.5 ± 11.3%; p = 0.0378; n = 4) compared with primary microglia. (H) CD11b was not altered in any condition. Each bar represents the mean ± SEM. * p<0.05; ** p<0.01; ^ns^ not significant. Neu = neurons; MG = microglia; Astro = astrocytes; ArgI = arginase I.

### 3.3. Astrocytes turn to a less reactive phenotype in the triple co-culture model

Once we observed that microglia were less inflammatory in the triple co-culture, we addressed astrocytic activation in the model. Besides, reactive microglia are necessary to induce A1 reactive astrocytes *in vitro* via secreted signaling molecules (Liddelow *et al*., 2017).

Firstly, we analyzed astrocytic morphology as a measure of their activation state. We used the astrocytic marker GFAP to obtain morphological parameters. We observed that, in the triple co-culture, astrocytes significantly increased the parameters of density (18.9 ± 3.7%; p = 0.0292; n = 4) and span ratio (24.9 ± 6.4%; p = 0.0401; n = 4), while they reduced their circularity (11.9 ± 2.3%; p = 0.0123; n = 4) compared with primary astrocytes (Figure 3A-D). These parameters in the triple co-culture reflect an increase in cellular ramifications, which resembles better the physiological resting state. The presence of neurons was enough to see a morphological change in astrocytes, although it was not statistically significant.

**Figure 3.**
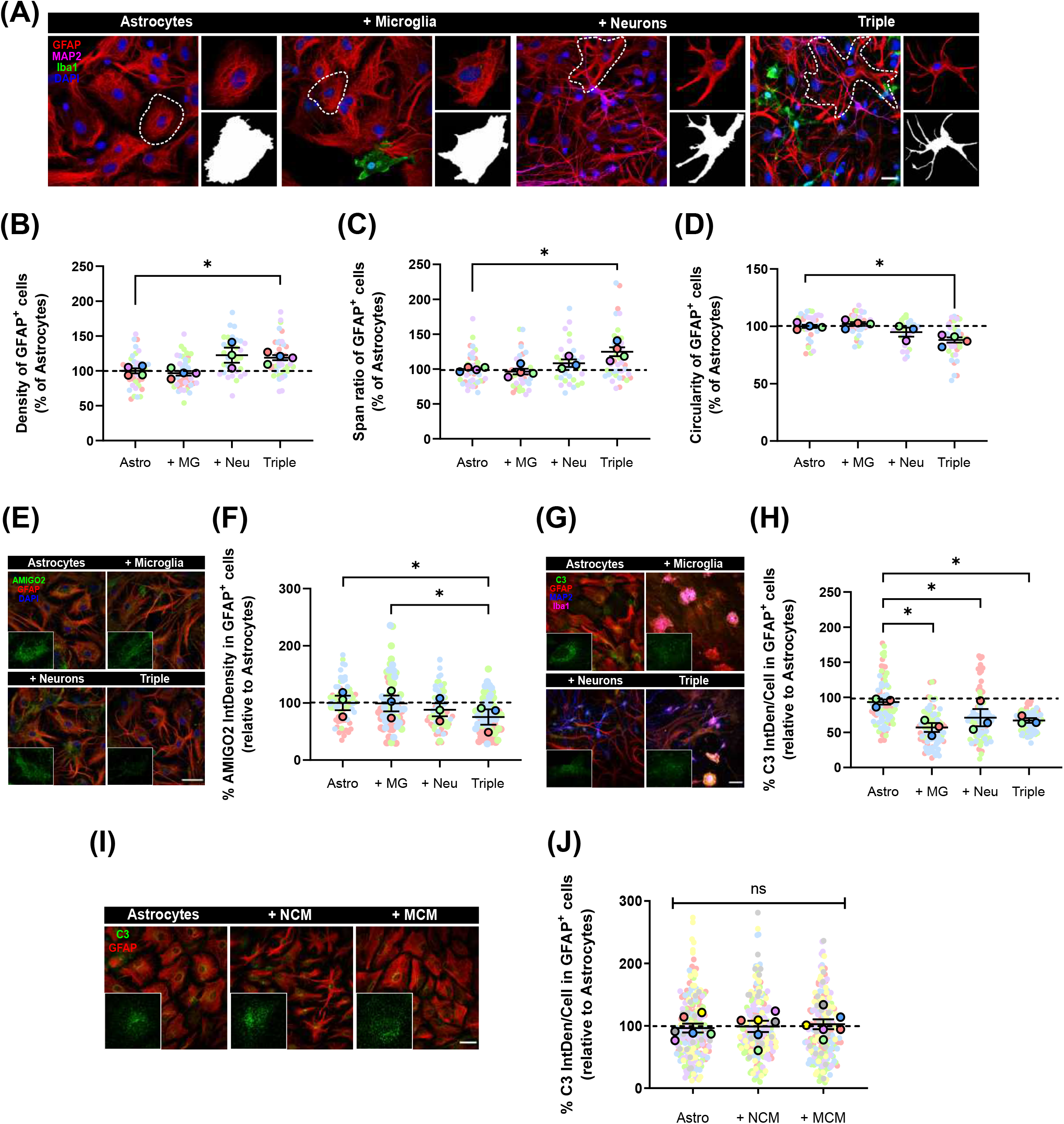
Astrocytes displayed a less reactive morphology and reduced expression of A1 activation markers AMIGO2 and C3. (A) Representative images of the quantification of astrocytic morphology using GFAP and Fractal Analysis in FIJI Software. Black and white images represent the outline of individual cells selected in each condition. Scale bar = 20 μm. (B) Astrocytes in the triple co-culture showed an increase in density (18.9 ± 3.7%; p = 0.0292; n = 4) and (C) span ratio (24.9 ± 6.4%; p = 0.0401; n = 4), while they displayed a decrease in (D) circularity (11.9 ± 2.3%; p = 0.0123; n = 4), compared with primary astrocytes alone. (E) Representative image of the quantification of A1 astrocytic marker AMIGO2. Scale bar = 40 μm. (F) Astrocytes decreased the expression of AMIGO2 in the triple co-culture compared with microglia-astrocyte co-culture (23.9 ± 13.4%; p = 0.0323; n = 3) and primary astrocytes (24.6 ± 13.4%; p = 0.0387; n = 3). (G) Representative image of the quantification of A1 astrocytic marker C3. Scale bar = 40 μm. (H) Expression of C3 decreased in the microglia-astrocyte co-culture (38.7 ± 6.4%; p = 0.0296; n = 3), in the neuronastrocyte co-culture (23.8 ± 12.3%; p = 0.0343; n = 3) and in the triple coculture (28.0 ± 3.3%; p = 0.0376; n = 3), compared with primary astrocytes. (I) Representative image of the quantification of C3 in GFAP^+^ astrocytes after conditioned media treatment. Scale bar = 40 μm. (J) C3 levels did not change with either NCM or MCM. Each bar represents the mean ± SEM. * p<0.05; ^ns^ not significant. Neu = neurons; MG = microglia; Astro = astrocytes; NCM = neuron conditioned media; MCM = microglia conditioned media.

We also quantified the expression of the two A1 astrocytic markers AMIGO2 and C3, using IF. In the triple co-culture, GFAP^+^ astrocytes significantly decreased the expression of AMIGO2 compared with microglia-astrocyte coculture (23.9 ± 13.4%; p = 0.0323; n = 3) and primary astrocytes (24.6 ± 13.4%; p = 0.0387; n = 3) (Figure 3E, F). In addition, C3 expression in astrocytes decreased in all of the co-culture conditions; in the microglia-astrocyte coculture (38.7 ± 6.4%; p = 0.0296; n = 3), in the neuron-astrocyte co-culture (23.8 ± 12.3%; p = 0.0343; n = 3) and in the triple co-culture (28.0 ± 3.3%; p = 0.0376; n = 3) (Figure 3G, H). In order to analyze if astrocytic C3 was modulated via cell-to-cell contact or via secreted factors, we used conditioned media. We incubated primary astrocytes with conditioned media from neurons (NCM) or from microglia (MCM) for three days, and then quantified C3. There were no changes in C3 levels with either NCM (p = 0.6454; n = 6) or MCM (p = 0.6897; n = 6), so physical contact between cells was required for astrocytes to reduce C3 expression (Figure 3I, J). In conclusion, astrocytes seemed less reactive in the triple co-culture environment, with a more ramified morphology and reduced expression of A1 markers AMIGO2 and C3.

### 3.4. Oligomeric Aβ induces synaptic loss and increases microglial activation in the triple co-culture

Once we characterized the triple co-culture, we used it to establish a model where we could study AD *in vitro*, without the limitations of single primary cultures. We used oAβ as the stimulus to induce neurodegeneration, since oligomers seem to be the most neurotoxic amyloid species (as reviewed in Glabe, 2006).

We treated our triple co-cultures with 3 μM oAβ for 24 h, and then analyzed neurodegeneration features such as synaptic loss and neuroinflammation. We quantified pre- and post-synaptic markers in the neuronal processes using IF and found that both Syn and Homer1 were reduced with the oAβ treatment (7.6 ± 8.1%; p = 0.0364; n = 5) and (17.5 ± 6.4%; p = 0.0263; n = 5) respectively, compared with controls (Figure 4A-D). Along with synapse loss, inflammation characterized by microglial CD11b increase is another key feature of Aβ-induced neurodegeneration (Rao *et al*., 2011). For that reason, we analyzed CD11b in total lysate samples with WB, as a measure of microglia-related inflammation. We observed that, while Iba1 expression did not change (p = 0.1968; n = 4), CD11b increased in microglia with the oAβ treatment (16.0 ± 14.3%; p = 0.0067; n = 4) compared with controls (Figure 4E-G). Besides, there was a trend to decrease of anti-inflammatory cytokine TGF-β1 levels in the triple co-culture, although not statistically significant (p = 0.3709; n = 4) (Figure 4H).

**Figure 4.**
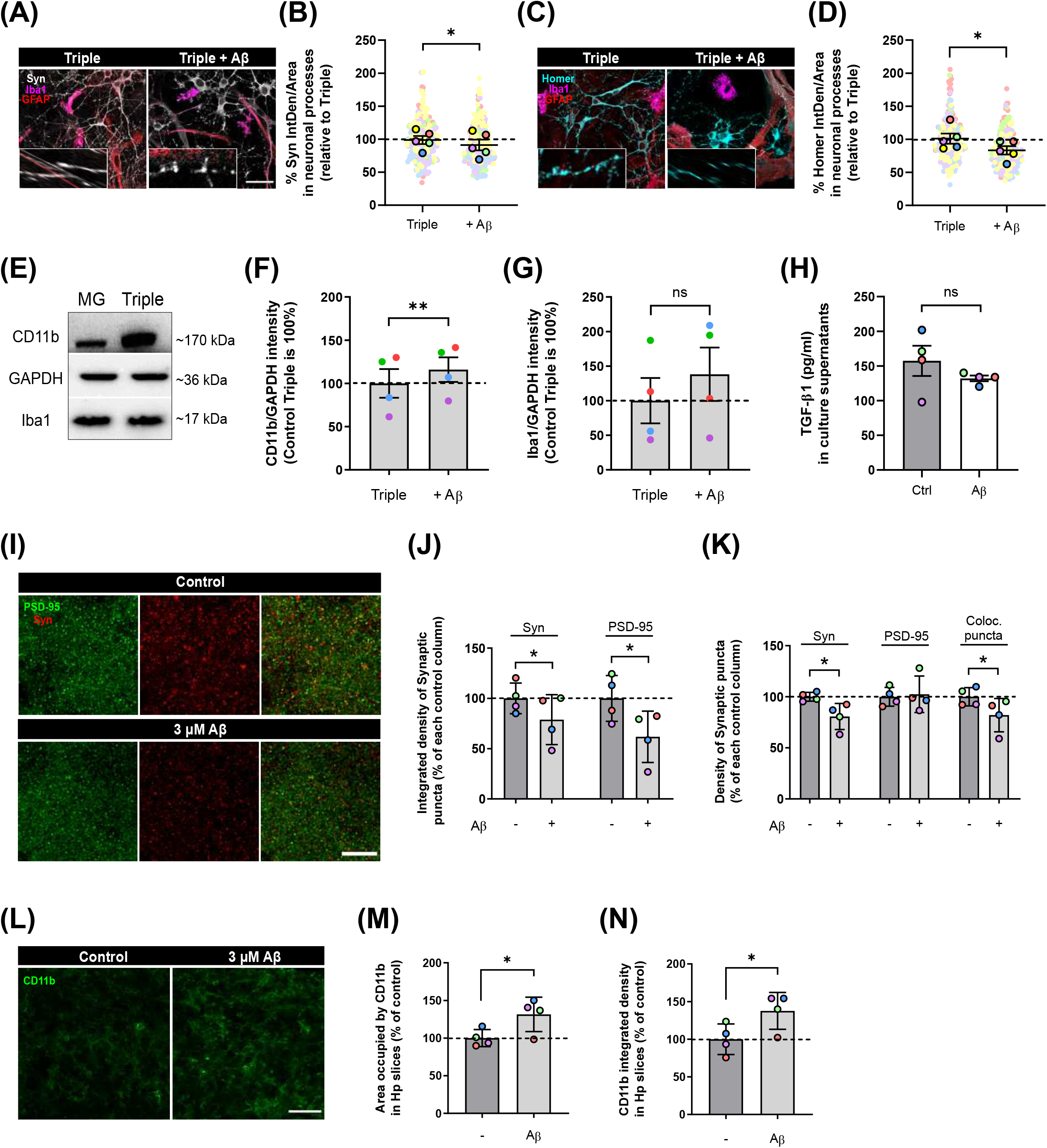
Oligomeric Aβ reduced pre- and post-synaptic puncta and increased microglial expression of CD11b in the triple co-culture and in hippocampal organotypic slices. (A) Synaptophysin in neuronal processes. Scale bar = 40 μm. (B) oAβ treatment reduced Synaptophysin (7.6 ± 8.1%; p = 0.0364; n = 5) in the triple co-culture. (C) Homer1 in neuronal processes. Scale bar = 40 μm. (D) Expression of Homer1 decreased with oAβ (17.5 ± 6.4%; p = 0.0263; n = 5). (E) Quantification of microglial markers in total lysates. (F) Activation marker CD11b increased in microglia with oAβ (16.0 ± 14.3%; p = 0.0067; n = 4) compared to controls. (G) Iba1 expression did not change with the treatment. (H) Cytokine TGF-β1 in culture supernatants. TGF-β1 levels tended to decrease in the triple co-culture with oAβ, although it was not statistically significant. (I) Quantification of synaptic markers in hippocampal organotypic slices. Scale bar = 10 μm. (J) Synaptophysin and PSD-95 integrated density were reduced with oAβ (21.1 ± 12.4%; p = 0.0470; n = 4) and (38.2 ± 12.8%; p = 0.0203; n = 4) respectively, compared to controls. (K) The density of Synaptophysin^+^ puncta was reduced (19.3 ± 6.4%; p = 0.0135; n = 4) with oAβ, along with the overall colocalization puncta (18.0 ± 8.1%; p = 0.0494; n = 4) with oAβ treatment. (L) CD11b staining in hippocampal organotypic slices. Scale bar = 40 μm. oAβ increased both the (M) area occupied by CD11b (31.5 ± 11.4%; p = 0.0309; n = 4) and its (N) integrated density (37.7 ± 12.3%; p = 0.0323; n = 4), compared to controls. Each bar represents the mean ± SEM. * p<0.05; ** p<0.01; ^ns^ not significant. Syn = synaptophysin.

To confirm our findings in the triple co-culture using a model with higher complexity, we used hippocampal organotypic cultures. We treated these organotypic cultures with the same conditions (3 μM oAβ for 24 h) and analyzed synaptic loss and neuroinflammation. Aβ treatment reduced the integrated density of Syn^+^ pre-synaptic puncta (21.1 ± 12.4%; p = 0.0470; n = 4) and PSD-95^+^ post-synaptic puncta (38.2 ± 12.8%; p = 0.0203; n = 4) compared with controls (Figure 4I, J). The puncta density was also decreased in the case of Syn (19.3 ± 6.4%; p = 0.0135; n = 4), which caused a reduction in the overall colocalization puncta (18.0 ± 8.1%; p = 0.0494; n = 4) (Figure 4K). Regarding microglial activation, we quantified CD11b^+^ microglia in the organotypic slices. We found that oAβ increased both the area occupied by CD11b (31.5 ± 11.4%; p = 0.0309; n = 4) and its integrated density (37.7 ± 12.3%; p = 0.0323; n = 4) compared to controls (Figure 4L-N). In conclusion, we were able to recapitulate Aβ-induced synaptic loss and microglial activation in our triple co-culture, and we confirmed these findings using hippocampal organotypic slices.

## 4. Discussion

We developed a murine triple co-culture model including neurons, microglia and astrocytes, that holds physiological characteristics that are lost in classical primary cultures. Microglia are less inflammatory, astrocytes are less reactive and neurons display a more mature morphology than in individual primary cultures. With this model, we were able to recapitulate Aβ-induced AD pathological features like synaptic loss and microglial activation, which will allow us to study neurodegeneration and neuroinflammation processes relevant to AD progression.

In our triple co-culture, neurons developed a higher number of post-synapses, while pre-synapses remained unchanged. This finding would align with previous work indicating that synaptogenesis promoted by astrocyte-secreted factors such as thrombospondins formed structurally normal synapses that are pre-synaptically active but post-synaptically silent (Christopherson *et al*., 2005). Our model would allow further functional studies of neuronal activity, as well as the identification of glial factors with potential synaptogenic activity. We also observed that the presence of both microglia and astrocytes in the culture promoted a significant morphological change in neurons, which developed a higher number of branches that were also longer. These findings, along with previous studies demonstrating that the presence of astrocytes increases neuronal viability *in vitro* (Aebersold *et al*., 2018), support our co-culture as a relevant system to study neuronal function in AD and other neurodegenerative diseases.

Microglial genomic and proteomic profile is very dependent on the environment. Previous studies have shown a differential gene expression in primary microglia as opposed to the adult CNS, revealed by a dedifferentiation of microglia in culture (Schmid *et al*., 2009). The use of serum as a way to stimulate cell proliferation and survival *in vitro* also causes relevant phenotypic changes in microglia. Bohlen *et al*. demonstrated that serum induced significant alterations in microglial morphology and intrinsic phagocytic capacity, and that these alterations could be avoided using serum-free astrocyte conditioned media containing CSF-1/IL-34, TGF-β2, and cholesterol (Bohlen *et al*., 2017). In the case of our triple co-culture model, we used serum-free neurobasal medium, which allowed us to avoid serum-related changes while maintaining neuronal viability. In these serum-free co-culture conditions, microglia decreased the expression of pro-inflammatory markers iNOS and IL-1β, while increased the expression of anti-inflammatory markers Arginase I and TGF-β1. We consider that both the serum-free conditions and the presence of neurons and astrocytes, contributed to maintain microglia in a less reactive state compared with primary cultures. Previous studies performed with a triple co-culture that included microglial cells N11, neuroblastoma N2A cells and brain microvascular endothelial MVEC(B3) cells in a transwell system shown that microglial stimulation with LPS induced inflammatory pathways that resulted in neuron apoptosis and damaged endothelial integrity (Zheng *et al*., 2021). Therefore, future studies including neuroinflammation induction (e.g. with LPS) would be required to have a more extensive phenotypic characterization of microglia in our model. Activated microglia also have a direct effect on astrocytes, inducing their shift to A1 reactive phenotype by the secretion of IL-1α, TNF and C1q (Liddelow *et al*., 2017). In our model, we observed that, microglia being less reactive, astrocytes had a ramified morphology and decreased the expression of A1 activation markers AMIGO2 and C3. On the contrary, we did not observe any expression changes in astrocytic C3 with conditioned media. These findings seem to contradict previous reports that, using a tri-culture system with hPSC-derived microglia, astrocytes, and neurons, found an increase in astrocytic C3 in the presence of microglia and microglia conditioned media (Guttikonda *et al*., 2021). This discrepancy could be because the authors quantified secreted C3, while we quantified C3 within astrocytes. However, a more exhaustive analysis will be necessary to understand the communication processes between cell types.

Lastly, we used oligomeric Aβ to model amyloid pathology. With this, we were able to observe pre- and post-synaptic loss, as well as a microglial activation reflected by an increase in marker CD11b. Both microglia and astrocytes can participate in synaptic elimination via the complement system (Hong *et al*., 2016) and MEGF10/MERTK pathways (Chung *et al*., 2013), respectively. Therefore, our model could be useful to further study the role of these pathways in Aβ-induced synaptic loss. We used a more complex model such as hippocampal organotypic cultures to confirm our findings and observed the same negative effects of oligomeric Aβ on synaptic loss and microglial activation. Compared with organotypic cultures, our triple co-culture model has a defined concentration of each cell type that can be manipulated, becoming a complementary model for mechanistic studies.

Altogether, our findings suggest that the neuron, microglia, and astrocyte triple co-culture model we developed is reliable, affordable and a useful tool for the study of mechanisms implicated in AD neurodegeneration. In agreement with previous reports with *in vitro* models, ours also closely mimics the *in vivo* environment, where the cross talk between neurons and glial cells contributes to neuroinflammation processes (Goshi *et al*., 2020), and where alterations in glial cells can lead to neurodegeneration (Lange *et al*., 2018). Our model can also be potentially used in high throughput screening setups, so we expect that it will help finding new therapeutic targets for AD and other neurodegenerative diseases.

## Acknowledgments

Authors acknowledge financial support by Basque Government (IT1203-19; ELKARTEK KK-2020/00034; PIBA_2020_1_0012), CIBERNED (CB06/0005/0076), MICINN (PID2019-109724RB-I00). CL and JZI were supported by PhD Scholarships from the Tatiana Pérez de Guzmán el Bueno Foundation and Basque Government respectively.

## Authors’ contributions

ECZ and CL conceived the experiments. CL, and JZI performed the experiments. JV provided tools for image analysis. ECZ and CL wrote the paper. EA and CM provided valuable insight and advice throughout the project. All authors read and approved the final manuscript.

## References

Aebersold MJ, Thompson-Steckel G, Joutang A, Schneider M, Burchert C, Forró C, et al. Simple and Inexpensive Paper-Based Astrocyte Co-culture to Improve Survival of Low-Density Neuronal Networks. Front Neurosci 2018; 0: 94.

Alberdi E, Sánchez-Gómez MV, Cavaliere F, Pérez-Samartín A, Zugaza JL, Trullas R, et al. Amyloid β oligomers induce Ca2+ dysregulation and neuronal death through activation of ionotropic glutamate receptors. Cell Calcium 2010; 47: 264–72.

Alberdi E, Wyssenbach A, Alberdi M, Sánchez-Gómez M V., Cavaliere F, Rodríguez JJ, et al. Ca ^2+^-dependent endoplasmic reticulum stress correlates with astrogliosis in oligomeric amyloid β-treated astrocytes and in a model of Alzheimer’s disease. Aging Cell 2013; 12: 292–302.

Bohlen CJ, Bennett FC, Tucker AF, Collins HY, Mulinyawe SB, Barres BA. Diverse Requirements for Microglial Survival, Specification, and Function Revealed by Defined-Medium Cultures. Neuron 2017; 94: 759–773.e8.

Brewer GJ, Torricelli JR, Evege EK, Price PJ. Optimized survival of hippocampal neurons in B27-supplemented neurobasal™, a new serum-free medium combination. J Neurosci Res 1993; 35: 567–76.

Castellani RJ, Rolston RK, Smith MA. Alzheimer Disease. Dis Mon 2010; 56: 484.

Christopherson KS, Ullian EM, Stokes CCA, Mullowney CE, Hell JW, Agah A, et al. Thrombospondins are astrocyte-secreted proteins that promote CNS synaptogenesis. Cell 2005; 120: 421–33.

Chung W-S, Clarke LE, Wang GX, Stafford BK, Sher A, Chakraborty C, et al. Astrocytes mediate synapse elimination through MEGF10 and MERTK pathways. Nature 2013; 504: 394–400.

Correa FG, Hernangómez M, Guaza C. Understanding microglia-neuron cross talk: Relevance of the microglia-neuron cocultures. Methods Mol Biol 2013; 1041: 215–29.

Czirr E, Castello NA, Mosher KI, Castellano JM, Hinkson I V., Lucin KM, et al. Microglial complement receptor 3 regulates brain Aβ levels through secreted proteolytic activity. J Exp Med 2017; 214: 1081–92.

Dahlgren KN, Manelli AM, Blaine Stine W, Baker LK, Krafft GA, Ladu MJ. Oligomeric and Fibrillar Species of Amyloid-β Peptides Differentially Affect Neuronal Viability *. J Biol Chem 2002; 277: 32046–53.

Du F, Yu Q, Chen A, Chen D, ShiDu Yan S. Astrocytes Attenuate Mitochondrial Dysfunctions in Human Dopaminergic Neurons Derived from iPSC. Stem Cell Reports 2018; 10: 366–74.

Floden AM, Li S, Combs CK. β-Amyloid-Stimulated Microglia Induce Neuron Death via Synergistic Stimulation of Tumor Necrosis Factor α and NMDA Receptors. J Neurosci 2005; 25: 2566–75.

Furman JL, Sama DM, Gant JC, Beckett TL, Murphy MP, Bachstetter AD, et al. Targeting Astrocytes Ameliorates Neurologic Changes in a Mouse Model of Alzheimer’s Disease. J Neurosci 2012; 32: 16129–40.

Glabe CG. Common mechanisms of amyloid oligomer pathogenesis in degenerative disease. Neurobiol Aging 2006; 27: 570–5.

Goshi N, Morgan RK, Lein PJ, Seker E. A primary neural cell culture model to study neuron, astrocyte, and microglia interactions in neuroinflammation. J Neuroinflammation 2020; 17: 155.

Gosselin D, Skola D, Coufal NG, Holtman IR, Schlachetzki JCM, Sajti E, et al. An environment-dependent transcriptional network specifies human microglia identity. Science 2017; 356: 1248–59.

Guttikonda SR, Sikkema L, Tchieu J, Saurat N, Walsh R, Harschnitz O, et al. Fully defined human pluripotent stem cell-derived microglia and tri-culture system model C3 production in Alzheimer’s disease. Nat Neurosci 2021; 24: 343.

Haenseler W, Sansom SN, Buchrieser J, Newey SE, Moore CS, Nicholls FJ, et al. A Highly Efficient Human Pluripotent Stem Cell Microglia Model Displays a Neuronal-Co-culture-Specific Expression Profile and Inflammatory Response. Stem Cell Reports 2017; 8: 1727–42.

Hernangómez M, Mestre L, Correa FG, Loría F, Mecha M, Iñigo PM, et al. CD200-CD200R1 interaction contributes to neuroprotective effects of anandamide on experimentally induced inflammation. Glia 2012; 60: 1437–50.

Hong S, Beja-Glasser VF, Nfonoyim BM, Frouin A, Li S, Ramakrishnan S, et al. Complement and microglia mediate early synapse loss in Alzheimer mouse models. Science (80-) 2016; 352: 712–6.

Ibarretxe G, Sánchez-Gómez MV, Campos-Esparza MR, Alberdi E, Matute C. Differential oxidative stress in oligodendrocytes and neurons after excitotoxic insults and protection by natural polyphenols. Glia 2006; 53: 201–11.

Lange J, Haslett LJ, Lloyd-Evans E, Pocock JM, Sands MS, Williams BP, et al. Compromised astrocyte function and survival negatively impact neurons in infantile neuronal ceroid lipofuscinosis. Acta Neuropathol Commun 2018; 6: 74.

Larm JA, Cheung NS, Beart PM. (S)-5-fluorowillardiine-mediated neurotoxicity in cultured murine cortical neurones occurs via AMPA and kainate receptors. Eur J Pharmacol 1996; 314: 249–54.

Liddelow SA, Guttenplan KA, Clarke LE, Bennett FC, Bohlen CJ, Schirmer L, et al. Neurotoxic reactive astrocytes are induced by activated microglia. Nat 2017 5417638 2017; 541: 481–7.

Majumdar A, Capetillo-Zarate E, Cruz D, Gouras GK, Maxfield FR. Degradation of Alzheimer’s amyloid fibrils by microglia requires delivery of ClC-7 to lysosomes. Mol Biol Cell 2011; 22: 1664–76.

McCarthy KD, De Vellis J. Alpha-adrenergic receptor modulation of beta-adrenergic, adenosine and prostaglandin E1 increased adenosine 3’:5’-cyclic monophosphate levels in primary cultures of glia. J Cyclic Nucleotide Res 1978; 4: 15–26.

McCarthy KD, De Vellis J. Preparation of separate astroglial and oligodendroglial cell cultures from rat cerebral tissue. J Cell Biol 1980; 85: 890–902.

McFarland KN, Ceballos C, Rosario A, Ladd T, Moore B, Golde G, et al. Microglia show differential transcriptomic response to Aβ peptide aggregates ex vivo and in vivo. Life Sci Alliance 2021; 4–7.

Ortiz-Sanz C, Gaminde-Blasco A, Valero J, Bakota L, Brandt R, Zugaza JL, et al. Early Effects of Aβ Oligomers on Dendritic Spine Dynamics and Arborization in Hippocampal Neurons. Front Synaptic Neurosci 2020; 12: 2.

Paresce DM, Chung H, Maxfield FR. Slow degradation of aggregates of the Alzheimer’s disease amyloid β-protein by microglial cells. J Biol Chem 1997; 272: 29390–7.

Park J, Wetzel I, Marriott I, Dréau D, D’Avanzo C, Kim DY, et al. A 3D human triculture system modeling neurodegeneration and neuroinflammation in Alzheimer’s disease. Nat Neurosci 2018 217 2018; 21: 941–51.

Rao JS, Rapoport SI, Kim HW. Altered neuroinflammatory, arachidonic acid cascade and synaptic markers in postmortem Alzheimer’s disease brain. Transl Psychiatry 2011 18 2011; 1: e31–e31.

Schmid CD, Melchior B, Masek K, Puntambekar SS, Danielson PE, Lo DD, et al. Differential gene expression in LPS/IFNγ activated microglia and macrophages: In vitro versus in vivo. In: Journal of Neurochemistry. John Wiley & Sons, Ltd; 2009. p. 117–25

Stoppini L, Buchs PA, Muller D. A simple method for organotypic cultures of nervous tissue. J Neurosci Methods 1991; 37: 173–82.

Yoon JJ, Nicholson LFB, Feng SX, Vis JC, Green CR. A novel method of organotypic brain slice culture using connexin-specific antisense oligodeoxynucleotides to improve neuronal survival. Brain Res 2010; 1353: 194–203.

Young K, Morrison H. Quantifying microglia morphology from photomicrographs of immunohistochemistry prepared tissue using imagej. J Vis Exp 2018; 2018: 57648.

Zhang SC, Fedoroff S. Neuron-microglia interactions in vitro. Acta Neuropathol 1996; 91: 385–95.

Zheng Y-F, Zhou X, Chang D, Bhuyan DJ, Zhang JP, Yu W-Z, et al. A novel triculture model for neuroinflammation. J Neurochem 2021; 156: 249–61.

